# Molecular genetic contributions to social deprivation and household income in UK Biobank (n = 112,151)

**DOI:** 10.1101/043000

**Authors:** W David Hill, Saskia P Hagenaars, Riccardo E Marioni, Sarah E Harris, David CM Liewald, Gail Davies, International Consortium for Blood Pressure, Andrew M McIntosh, Catharine R Gale, Ian J Deary

## Abstract

Individuals with lower socio-economic status (SES) are at increased risk of physical and mental illnesses and tend to die at an earlier age [1-3]. Explanations for the association between SES and health typically focus on factors that are environmental in origin [4]. However, common single nucleotide polymorphisms (SNPs) have been found collectively to explain around 18% (SE = 5%) of the phenotypic variance of an area-based social deprivation measure of SES [5]. Molecular genetic studies have also shown that physical and psychiatric diseases are at least partly heritable [6]. It is possible, therefore, that phenotypic associations between SES and health arise partly due to a shared genetic etiology. We conducted a genome-wide association study (GWAS) on social deprivation and on household income using the 112,151 participants of UK Biobank. We find that common SNPs explain 21% (SE = 0.5%) of the variation in social deprivation and 11% (SE = 0.7%) in household income. Two independent SNPs attained genome-wide significance for household income, rs187848990 on chromosome 2, and rs8100891 on chromosome 19. Genes in the regions of these SNPs have been associated with intellectual disabilities, schizophrenia, and synaptic plasticity. Extensive genetic correlations were found between both measures of socioeconomic status and illnesses, anthropometric variables, psychiatric disorders, and cognitive ability. These findings show that some SNPs associated with SES are involved in the brain and central nervous system. The genetic associations with SES are probably mediated via other partly-heritable variables, including cognitive ability, education, personality, and health.

## Results and Discussion

Using GCTA-GREML [7], we first estimated the heritability of each of the SES variables and found that a total of 21% (SE = 0.5%) of phenotypic variation in social deprivation and 11% (SE= 0.7%) of household income was explained by the additive effects of common SNPs. Next, genome-wide association analyses for social deprivation and household income were performed using an imputed dataset that combined the UK10K haplotype and 1000 Genomes Phase 3 reference panels; details can be found at http://biobank.ctsu.ox.ac.uk/crystal/refer.cgi?id=157020. We found no genome-wide significant findings associated with social deprivation (See Figure 1 and Figure S1). For household income, four SNPs attained genome-wide significance (*P* < 5 × 10^-8^), rs187848990 on chromosome 2, and rs7252896 rs7255223 and rs8100891 on chromosome 19 (See Figure 2, Figure S1 and Table S1).

**Figure 1.**
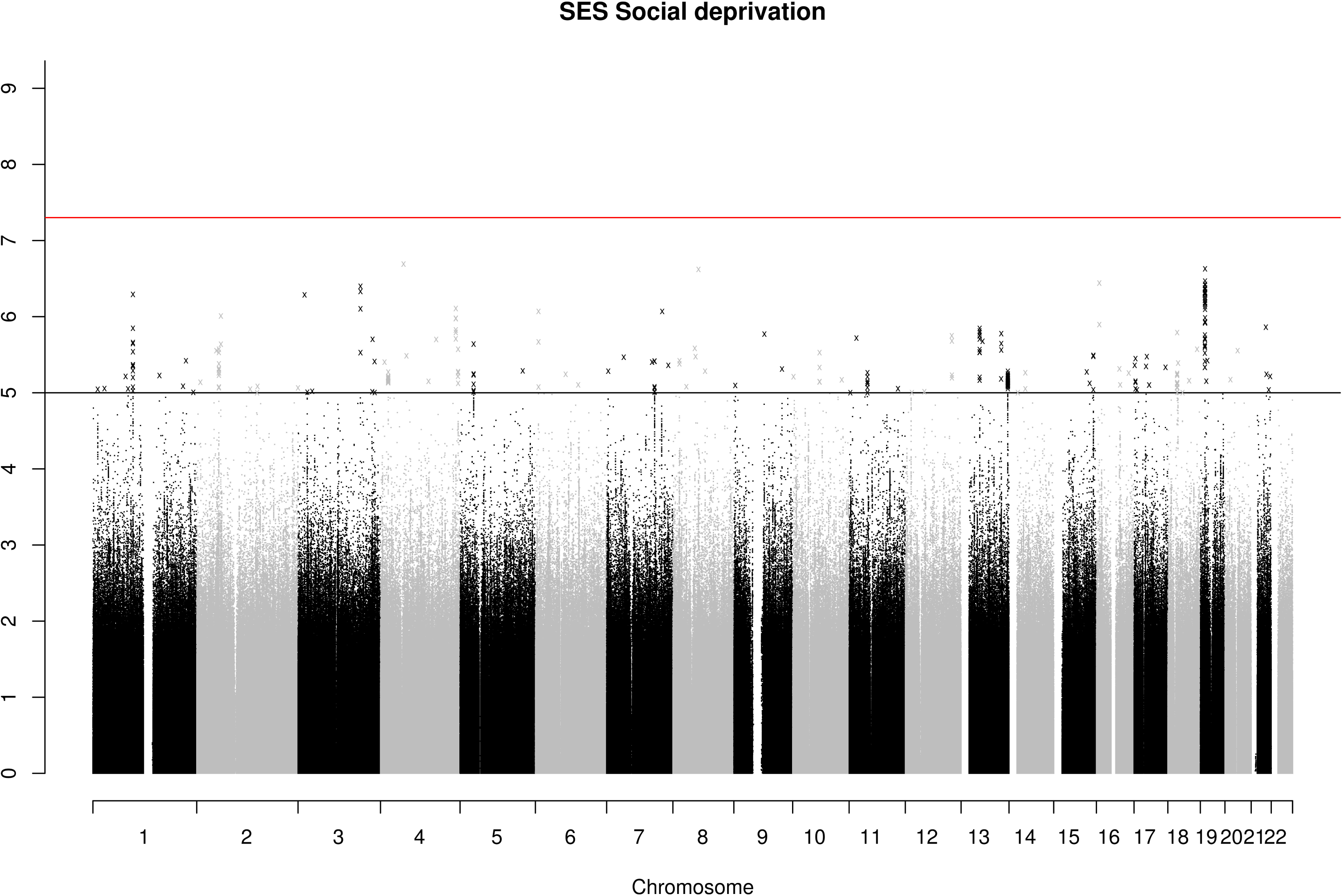
Manhattan plot of ‒log10 (*P* values) for social deprivation. The red line indicates genome-wide significance (*P* < 5x10^-8^). The black line indicates values that were suggestive of statistical significance (*P* < 1 × 10^-5^).

**Figure 2.**
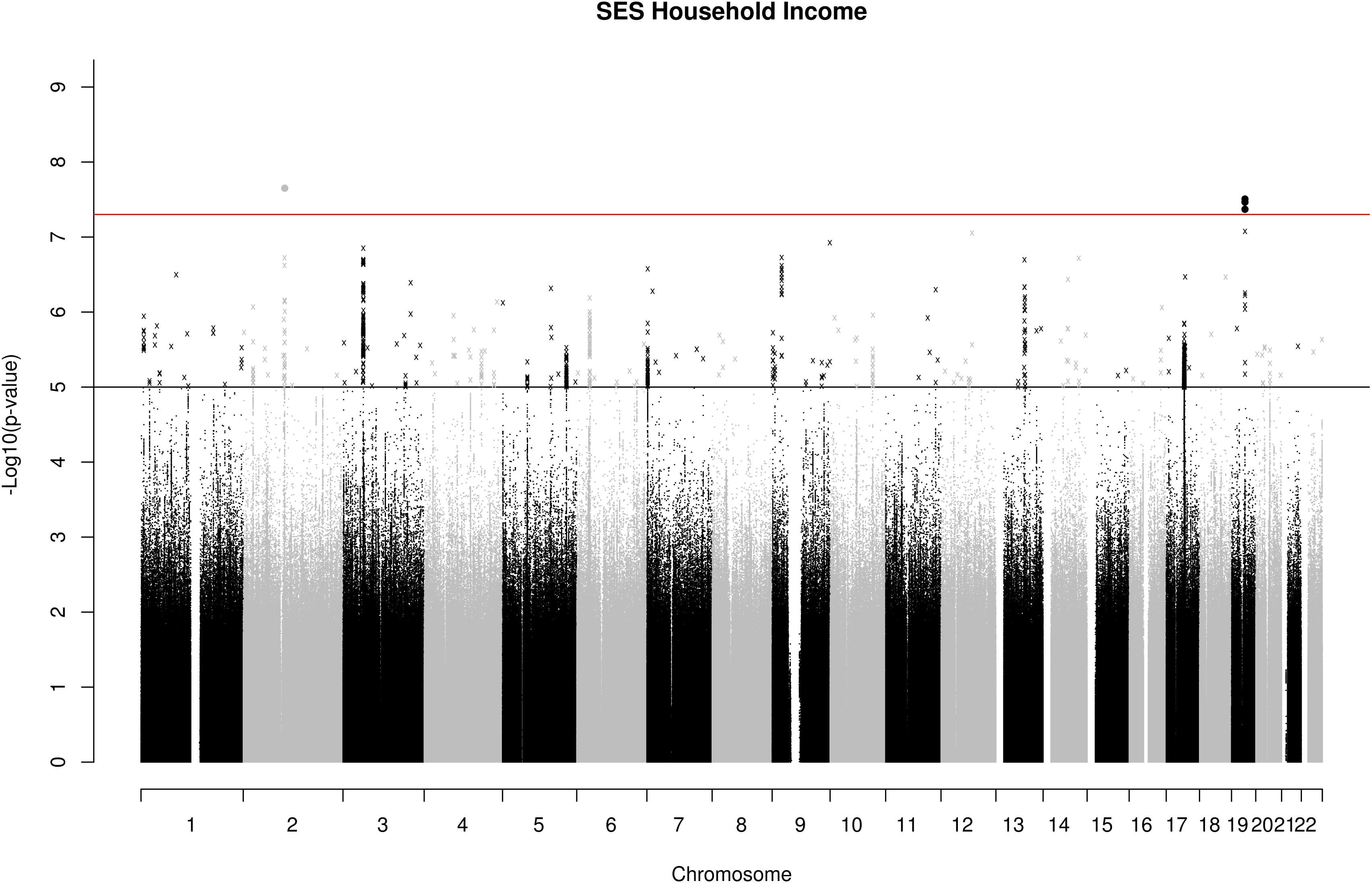
Manhattan plot of ‒log10 (*P* values) for household income. The red line indicates genome wide significance (*P* < 5x10^-8^). The black line indicates values that were suggestive of statistical significance (*P* < 1 x 10^-5^).

The “clump” function in PLINK [8] was used to identify patterns of linkage disequilibrium in the data set and showed that these four SNPs were located in two independent regions (Figure S2). The region on chromosome 2 spanned 583kb and the most significant SNP was rs187848990, *P* = 2.325x10^-8^. This region contains five genes: *AFF3*, *CHST10*, *LONRF2*, *NMS*, and *PDCL3.* The *AFF3* gene has previously been associated with intellectual disability [9], and *CHST10* is involved with synaptic plasticity [10]. The region on chromosome 19 spanned 18kb with the most significant SNP being rs8100891, *P* = 3.423 × 10^-8^. This region contains the gene *ZNF507* which has been implicated in neurodevelopmental disorders including schizophrenia [11], a disorder affecting cognitive function, that shows a strong genetic correlation with intelligence [12]. It is possible, therefore, that these genetic associations with SES may be mediated, in part, through cognitive ability; it is well established that an individual’s level of cognitive ability is predictive of their socioeconomic status [13].

Using the GTEx database (http://www.broadinstitute.org/gtex/), cis-eQTL associations were identified for the four household income genome-wide significant SNPs (Table S2). For this study, data mining of regulatory elements was restricted to normal tissues. There was evidence of regulatory elements associated with all four of the genome-wide significant SNPs (Table S2).

Gene based association testing was conducted using MAGMA [14]. Following Bonferroni correction for multiple testing (*α* = 2.768 × 10^-6^), gene based association tests identified one gene associated with social deprivation, *ACCSL* on chromosome 11. For household income 12 genes showed a significant association: *KANSL1*, *MST1*, *RNF123*, *MAPT*, *APEH*, *BSN*, *PLEKHM1*, *SGCD*, *DAG1*, *CRHR1*, *AMT*, and *ZDHHC11* (Supplemental Experimental Procedures). The *MAPT*,*KANSL1*, *PLEKHM1*, and *CRHR1* genes have been associated with Alzheimer’s disease [15]. Tables S3 and S4 show the top 20 genes and their level of association with social deprivation and household income.

Partitioned heritability analysis was then conducted on both SES phenotypes [16]. The goal of the partitioned heritability analysis was to determine if SNPs grouped together, according to a specific biological function or role, make an enriched contribution to the total proportion of heritability for each of the SES variables. The functional categories used here overlap considerably, meaning that the heritability measured by all groups, when they are summed, can exceed 100%. Figures S3 and S4 show the results of the enrichment analysis for social deprivation and for household income. By deriving a heritability estimate for functional classes of SNPs across the genome, a significant enrichment was found for evolutionarily conserved regions of the genome. These evolutionarily conserved regions accounted for 2.6% of the SNPs in both SES phenotypes and accounted for 44% (SE = 12%) of the heritability of social deprivation and 53% (SE = 12%) of the heritability for household income. The trend for an enrichment in heritability emanating from such regions is consistent with the results from other quantitative traits and diseases [16] and, as such, further highlights the importance of these genetic regions as sources of phenotypic variance across traits. Under models of neutral selective pressure, these regions accumulate base-pair substitutions at a lower rate, indicative of their being regions where mutation results in the production of phenotypic variance susceptible to the effects of natural selection. Genetic variance within these regions may highlight a role for disease-causing loci which in turn might account for some phenotypic variance in SES. However, it is also possible that, as intelligence is phenotypically and genetically associated with many health traits [17] and thought to be evolutionarily selected for [18], these regions may show their association with SES partly through cognitive differences. These two explanations are not mutually exclusive because, after intelligence is included as a covariate, the associations between adult SES and health outcomes, although attenuated, remain significant [1].

The functional category of H3K9ac also showed a statistically significant enrichment for household income. This grouping contained 23% of the SNPs which collectively explained 64% (SE = 11%) of its SNP-based heritability. This functional group corresponds to histone marks, which are post-translational alterations of histones that operate either to facilitate or to repress the level of gene expression. Specifically, H3K9ac pertains to the acetylation of histone H3 at lysine 9. The Enhancer regions and DNase I hypersensitivity sites (DHS) also showed significant enrichment for household income. Enhancer regions included 15% of the SNPs which collectively explained 48% (SE = 12%) of the SNP-based heritability. The DHS functional category contained 50% of the SNPs collectively accounting for 100% of the heritability estimate (SE = 17%).

A cell-specific analysis of 10 broad tissue types was conducted (See Supplementary Experimental Information). Figures S5 and S6 show the results of cell specific enrichment for social deprivation and household income. For social deprivation, significant enrichment was found in variants exerting an effect within the central nervous system. Variants expressed in the central nervous system accounted for 15% of the total number of SNPs, but accounted for 47% (SE = 11%) of the heritability of social deprivation and 37% (SE = 9%) of the heritability of household income. For household income, this did not survive multiple testing correction.

We next derived genetic correlations, using linkage disequilibrium score regression (LDS regression)[19], between both measures of SES and a set of 32 phenotypes that have all been shown to be phenotypically associated with SES. Table S5 provides a reference describing the phenotypic associations between measures of SES and broadly-conceived health variables. Full details of the GWAS that provided summary statistics for each of the 32 phenotypes, along with links to the data, are provided in Table S6. The direction of effect for the genetic correlations and polygenic profile scores examining Townsend scores was reversed to facilitate a comparison with the household income variable.

Following false discovery rate (FDR) correction for multiple comparisons, 16 of the 34 genetic correlations were statistically significant for the Townsend social deprivation measure (See Figure 3 & Tables S7), and 24 of the 34 were significant for household income (Figure 4 & Table S8). The large number of genetic correlations found indicates that the molecular genetic associations with SES overlap with many other phenotypes. A large degree of overlap was found for variables that are cognitive in nature. Significant genetic correlations were observed, for example, between both measures of SES and childhood cognitive ability (social deprivation, *r*_g_ = 0.500, income, *r*_g_ = 0.668), and also with longevity (social deprivation, *r*_g_ = 0.301, household income, *r*_g_ = 0.303). The direction of effect in each instance indicates that more affluent SES is associated with longer life and higher childhood intelligence. The average age of the participants in the GWAS for childhood intelligence was 11 years, whereas the measurements of SES from UK Biobank were taken at a mean age of 57 years. The finding of a genetic correlation between these two traits may indicate that a set of genetic variants contribute to higher intelligence, which in turn contributes to a higher SES in midlife. Significant genetic correlations were found between household income and intracranial volume and infant head circumference (*r*_g_ = 0.533, *r*_g_ = 0.239, respectively), indicating that the variants associated with brain size may also be associated with the amount of money brought into a household. The genetic correlation of 0.34, (SE = 0.07) between verbal numerical reasoning and the Townsend social deprivation index in UK Biobank was reported by us elsewhere (Marioni et al., in preparation).

**Figure 3.**
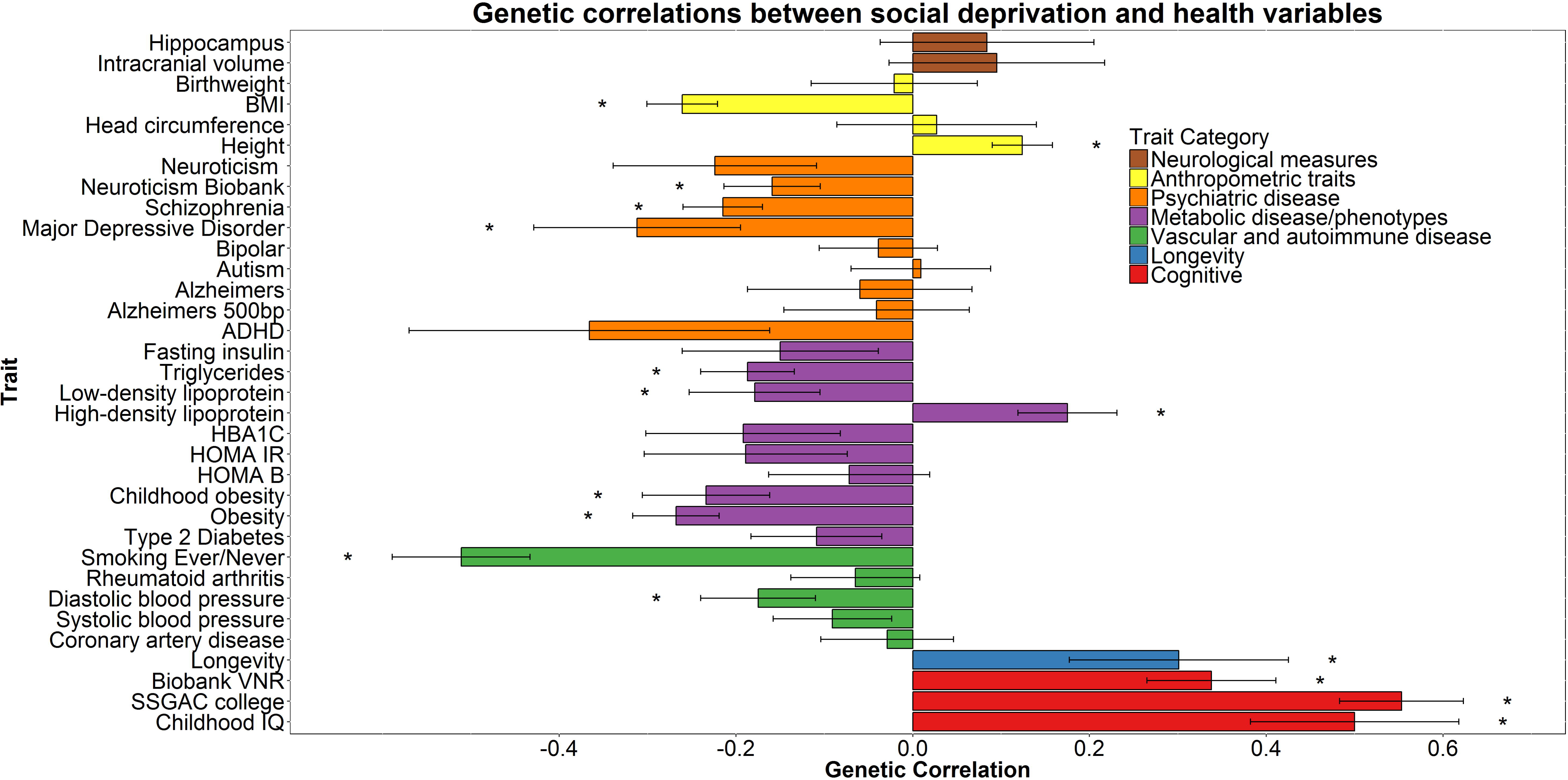
Genetic correlations between social deprivation (scores were reversed so that a higher Townsend score indicates a higher SES) and health and anthropometric variables. The x-axis depicts the magnitude of the genetic correlations and the y axis shows each trait. Statistical significance is indicated by asterisk. Statistical significance at *P* = 0.01535. Abbreviations: SSGAC, social science genetic associations consortium; HOMA B, homeostatic model assessment beta-cells; HOMA IR, homeostatic model assessment insulin resistance; HBA1C, glycated hemoglobin; ADHD, attention deficit hyperactivity disorder; MDD, major depressive disorder; BMI, body mass index; ICV, intracranial volume.

**Figure 4.**
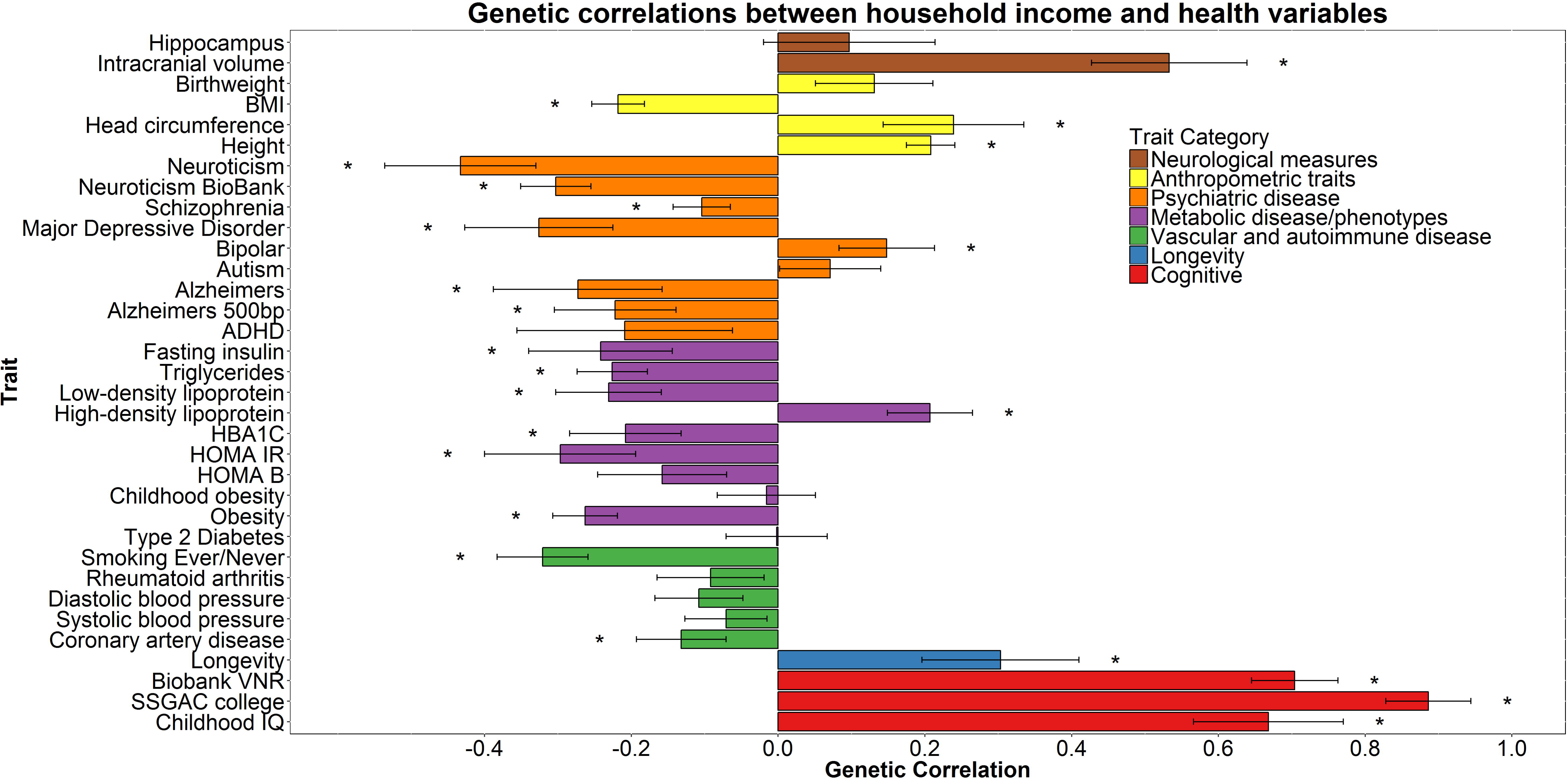
Genetic correlations between household income (where higher scores represent higher income) and health and anthropometric variables. The x-axis depicts the magnitude of the genetic correlations and the y axis shows each trait. Statistical significance is indicated by asterisk. Statistical significance at *P* = 0.032. Abbreviations: SSGAC, social science genetic associations consortium; HOMA B, homeostatic model assessment beta-cells; HOMA IR, homeostatic model assessment insulin resistance; HBA1C, glycated hemoglobin; ADHD, attention deficit hyperactivity disorder; MDD, major depressive disorder; BMI, body mass index; ICV, intracranial volume.

A noteworthy feature of our findings is that the pattern of genetic correlations between our two measures of SES‐‐one area-based and one individual-based‐‐was very similar and the genetic correlation between the two measures of SES was 0.871, (SE = 0.064). The Townsend social deprivation score is widely used as a proxy indicator of adult socioeconomic status, usually in the absence of an individual measure. It has been shown to be predictive of cancer incidence, all-cause mortality and other health outcomes [20]. Such area-level effects may comprise both compositional effects, i.e. effects that can be explained in terms of the characteristics of the residents of those areas, and contextual effects, i.e. effects that can be explained in terms of the characteristics of the areas. Although ecological correlations cannot be used to make causal inferences about individuals—the ecological fallacy—it has been argued that they arise largely from associations at the individual level [21]. There is evidence that area-based Townsend scores correlate highly with a similar measure of deprivation calculated at the individual level [22]. In our UK Biobank sample, where the individual-level measure of SES was based on household income alone, its correlation with Townsend score was small to moderate in size (r = 0.24); however, despite this, the pattern of genetic correlations between these two measures was very similar.

There are at least two explanations for the genetic SES-health correlations found using the LDS regression method. The first is that the genetic correlations might have been found as a result of the same genetic variants being directly involved in two phenotypes. The second is the notion of mediated pleiotropy, which describes situations where a phenotype is causally related to another; therefore, if a genetic variant is associated with the first phenotype it will be indirectly associated with the second [23]. Should multiple variants be used to establish pleiotropy, such as when using the LDS regression method, both of these forms of pleiotropy may apply, at different loci. However, because SES has no clear biological analog—it describes the environment of an individual or their achievements in it—mediated pleiotropy thorough intelligence or personality traits such as conscientiousness, for example, would appear to be a much more likely interpretation than biological pleiotropy.

The associations between rs187848990 and rs8100891 with household income, along with the heritability estimates for both measures of SES, are also most likely to be the result of mediation through other phenotypes. As genetic differences will not directly result in individual differences in SES, they may be contributing to differences in intelligence, personality, resistance to diseases, and other factors which, in turn, can contribute to differences in SES.

The effect sizes found for each individual SNP were small; however, as for any other polygenic trait, it is the combined effect of multiple SNPs that gives rise to observable phenotypic variance. Polygenic profile scores were created for 28 health-related phenotypes using published GWAS in all participants with genome-wide SNP data. When predicting into social deprivation based on the Townsend score, a total of 10 polygenic profile scores derived from GWAS summary data could predict a significant proportion of phenotypic variance (Table S9 for the most predictive model and Table S10 for results across *P* value thresholds). When predicting into household income, 27 out of 28 demonstrated statistical significance (Table S11 for the most predictive model and Table S12 for results across *P* value thresholds). Polygenic profile scores explained only a very small proportion of variance, but they illustrate that genetic risk for a range of diseases and cognitive ability can predict variance in these two SES measures.

The results here show that 21% of individual differences in area level social deprivation and 11% of household income can be explained by additive common genetic factors. Four genome-wide significant SNPs were found for household income, leading to the identification of two independent genomic regions containing genes with known associations with intellectual disabilities, synaptic plasticity, and schizophrenia. Extensive genetic correlations were found between both measures of SES and health-related traits, indicating a highly diffuse genetic architecture. These genetic correlations might provide a partial explanation for the phenotypic association between SES and health the majority of which, we think, is due to environmental factors.

## Experimental Procedure

Data were used from UK Biobank (http://www.ukbiobank.ac.uk) [24] to examine two measures of SES, the Townsend Social Deprivation Index [25], a measure of the social deprivation of the area in which the participant lives, and household income. A total of 112,005 individuals had a Townsend score, of whom 52.53% were female (Mean age = 56.91 years, SD = 7.93, range 40-73). A total of 96,900 participants had data pertaining to household income of whom 50.64% were female (Mean age = 56.53 years, SD = 7.95, range 40-73). Participants had undergone genome-wide SNP genotyping.

We conducted separate analyses for the Townsend deprivation score and household income. All analyses included only those individuals who described themselves as white British. All phenotypes were adjusted for age, gender, assessment center, genotyping batch, genotyping array, and 10 principal components in order to correct for population stratification prior to all analyses. See Supplemental Experimental Information for full description of genotyping, imputation, and the phenotypes used.

## Author contributions

Conceptualization, WDH, CRG, IJD.; Software DCML, GD.; Formal Analysis, WDH,SPH,REM.; Investigation, WDH.; Resources, International Consortium for Blood Pressure.; Data curation, WDH, SPH, SEH.; Writing – original draft, WDH. Writing Review and Editing, WDH, SPH, REM, SEH, DCML, GD, International Consortium for Blood Pressure, AMM CRG, IJD.

## Acknowledgements

This work was undertaken in The University of Edinburgh Centre for Cognitive Ageing and Cognitive Epidemiology (CCACE), supported by the cross-council Lifelong Health and Wellbeing initiative (MR/K026992/1). Funding from the Biotechnology and Biological Sciences Research Council (BBSRC), the Medical Research Council (MRC), and the University of Edinburgh and gratefully acknowledged. CCACE funding supports IJD.

WDH is supported by a grant from Age UK (Disconnected Mind Project).

We thank Stuart J Ritchie for comments on a draft of the manuscript.

## Declaration of interests

IJD is a participant in UK Biobank.

## References

1. Batty, G.D., Der, G., Macintyre, S., and Deary, I.J. (2006). Does IQ explain socioeconomic inequalities in health? Evidence from a population based cohort study in the west of Scotland. Bmj 332, 580–584.

2. Calixto, O.-J., and Anaya, J.-M. (2014). Socioeconomic status. The relationship with health and autoimmune diseases. Autoimmun. Rev. 13, 641–654.

3. Marmot, M.G., Stansfeld, S., Patel, C., North, F., Head, J., White, I., Brunner, E., Feeney, A., and Smith, G.D. (1991). Health inequalities among British civil servants: the Whitehall II study. The Lancet 337, 1387–1393.

4. Wilkinson, R.G., and Marmot, M.G. (2003). Social determinants of health: the solid facts, (Copenhagen).

5. Marioni, R.E., Davies, G., Hayward, C., Liewald, D., Kerr, S.M., Campbell, A., Luciano, M., Smith, B.H., Padmanabhan, S., Hocking, L.J., et al. (2014). Molecular genetic contributions to socioeconomic status and intelligence. Intelligence 44, 26–32.

6. Polderman, T.J., Benyamin, B., de Leeuw, C.A., Sullivan, P.F., van Bochoven, A., Visscher, P.M., and Posthuma, D. (2015). Meta-analysis of the heritability of human traits based on fifty years of twin studies. Nat. Genet. 47, 702–709.

7. Yang, J., Lee, S.H., Goddard, M.E., and Visscher, P.M. (2011). GCTA: a tool for genome-wide complex trait analysis. The American Journal of Human Genetics 88, 7.

8. Purcell, S., Neale, B., Todd-Brown, K., Thomas, L., Ferreira, M.A., Bender, D., Maller, J., Sklar, P., de Bakker, P.I., Daly, M.J., et al. (2007). PLINK: a tool set for whole-genome association and population-based linkage analyses. Am. J. Hum. Genet. 81, 559–575.

9. Melko, M., Douguet, D., Bensaid, M., Zongaro, S., Verheggen, C., Gecz, J., and Bardoni, B. (2011). Functional characterization of the AFF (AF4/FMR2) family of RNA-binding proteins: insights into the molecular pathology of FRAXE intellectual disability. Human molecular genetics 20, 1873–1885.

10. Ong, E., Yeh, J.C., Ding, Y., Hindsgaul, O., and Fukuda, M. (1998). Expression cloning of a human sulfotransferase that directs the synthesis of the HNK-1 glycan on the neural cell adhesion molecule and glycolipids. Journal of Biological Chemistry 273, 5190–5195.

11. Talkowski, M.E., Rosenfeld, J.A., Blumenthal, I., Pillalamarri, V., Chiang, C., Heilbut, A., Ernst, C., Hanscom, C., Rossin, E., Lindgren, A.M., et al. (2012). Sequencing chromosomal abnormalities reveals neurodevelopmental loci that confer risk across diagnostic boundaries. Cell 149, 525–537.

12. Hill, W.D., Davies, G., Liewald, D.C., McIntosh, A.M., Deary, I.J., and CHARGE Cognitive Working Group (2015). Age-dependent pleiotropy between general cognitive function and major psychiatric disorders. Biological psychiatry.

13. Strenze, T. (2007). Intelligence and socioeconomic success: A meta-analytic review of longitudinal research. Intelligence 35, 401–426.

14. de Leeuw, C.A., Mooij, J.M., Heskes, T., and Posthuma, D. (2015). MAGMA: Generalized Gene-Set Analysis of GWAS Data. PLoS Comp. Biol. 11.

15. Jun, G., Ibrahim-Verbaas, C., Vronskaya, M., Lambert, J., Chung, J., Naj, A., Kunkle, B., Wang, L., Bis, J., Bellenguez, C., et al. (2015). A novel Alzheimer disease locus located near the gene encoding tau protein. Mol. Psychiatry.

16. Finucane, H.K., Bulik-Sullivan, B., Gusev, A., Trynka, G., Reshef, Y., Loh, P.-R., Anttila, V., Xu, H., Zang, C., Farh, K., et al. (2015). Partitioning heritability by functional annotation using genome-wide association summary statistics. Nat. Genet. 47, 1228–1235.

17. Hagenaars, S.P., Harris, S.E., Davies, G., Hill, W.D., Liewald, D.C., Ritchie, S.J., Marioni, R.E., Fawns-Ritchie, C., Cullen, B., Malik, R., et al. (2015). Shared genetic aetiology between cognitive functions and physical and mental health in UK Biobank (N= 112 151) and 24 GWAS consortia. Mol. Psychiatry.

18. Penke, L., Denissen, J.J., and Miller, G.F. (2007). The evolutionary genetics of personality. European Journal of Personality 21, 549–587.

19. Bulik-Sullivan, B., Finucane, H.K., Anttila, V., Gusev, A., Day, F.R., Loh, P.-R., Duncan, L., Perry, J.R., Patterson, N., Robinson, E.B., et al. (2015). An atlas of genetic correlations across human diseases and traits. Nat. Genet.

20. Smith, G.D., Whitley, E., Dorling, D., and Gunnell, D. (2001). Area based measures of social and economic circumstances: cause specific mortality patterns depend on the choice of index. Journal of Epidemiology and Community Health 55, 149–150.

21. MacRae, K. (1994). Socioeconomic deprivation and health and the ecological fallacy. British Medical Journal 309, 1478–1479.

22. Adams, J., Ryan, V., and White, M. (2005). How accurate are Townsend Deprivation Scores as predictors of self-reported health? A comparison with individual level data. Journal of Public Health 27, 101–106.

23. Solovieff, N., Cotsapas, C., Lee, P.H., Purcell, S.M., and Smoller, J.W. (2013). Pleiotropy in complex traits: challenges and strategies. Nature Reviews Genetics 14, 483–495.

24. Sudlow, C., Gallacher, J., Allen, N., Beral, V., Burton, P., Danesh, J., Downey, P., Elliott, P., Green, J., Landray, M., et al. (2015). UK biobank: an open access resource for identifying the causes of a wide range of complex diseases of middle and old age. PLoS Med 12, e1001779.

25. Townsend, P. (1987). Deprivation. Journal of social policy. 16 2.

